# DID Neanderthals and Denisovans have *our* de novo genes?

**DOI:** 10.1101/003004

**Authors:** John S. Taylor

**Affiliations:** Department of Biology, University of Victoria, PO Box 1700, Station CSC, Victoria, BC, V8W 2Y2 Canada

## Abstract

Gene duplication provides a profusion of raw material for evolutionary innovation (Lynch and Conery, 2000). While most duplicates rapidly become unrecognizable some, e.g., those that are immediately useful or those that after a period of relaxed selection gain unique roles, are retained and thereby expand a genomes protein-coding repertoire. Ohno (1970) famously remarked that without gene duplication, the creation of metazoans, vertebrates, and mammals from unicellular organisms would have been impossible. Such big leaps in evolution, he argued, required the creation of new gene loci with previously nonexistent functions (Taylor and Raes, 2004). Recently, another source of new genes has been recognized: Though rare, it seems clear that new genes can emerge from formerly non-coding DNA, the ‘de novo’ protein coding genes (Zhao *et al.*, 2014, and references therein).

In 2009 Knowles and McLysaght reported the discovery of three human genes derived from non-coding DNA. They provided evidence that these genes, *CLUU1*, *C22orf45*, and *DNAH10OS*, were transcribed and translated, they identified orthologous non-coding DNA in chimpanzee (*Pan troglodytes*) and macaque (*Macaca mulatta*), and for each gene they located the critical ‘enabler’ mutations that extended the open reading frames (ORFs) allowing the production of a protein. These genes had no BLASTp hits in any other genome and were considered to be novel human genes, possibly responsible for human-specific traits.

Since the discovery of these genes, new high quality Denisovan and Neanderthal genomes have been reported. I used these resources in an effort to determine whether or not *CLUU1, C22orf45*, and *DNAH10OS* were truly human-specific. The Denisova Cave in the Altai Mountains of southern Siberia was the source of the bones for the Neanderthal and Denisovan genomes (Prüfer et al., 2013; Reich *et al.*, 2010; Meyer et al., 2012). Both bones were estimated to be between 30,000 and 50,000 years old. For the Neanderthal, bam files (which contain reads mapped to the human genome) for the de novo gene-bearing chromosomes (see below) were downloaded from the Max Plank Gesellschaft Department of Evolutionary Genetics (http://cdna.eva.mpg.de/neandertal/altai/bam/) and were compared to the human reference sequence (HG19) using the tview command in SAMtools (Li *et al.*, 2009). Denisovan sequence data were surveyed using the UCSC Genome Browser (Kent et al., 2002). All three de novo genes have a single ORF. *CLUU1* and *DNAH10OS* occur on human chromosome 12. For *CLUU1* the coding sequence begins at position 92818457 and ends at position 92818822 (HG19). *DNAH10OS* is encoded on the negative strand, beginning at position 124419130 and ending (upstream) at position 124418639 (HG19). *C22orf45* occurs on human chromosome 22, also on the negative strand: 22:24827520-24827041 (HG19).

For *CLUU1*, the enabler mutation is a single base (adenine) deletion (ΔA). This derived mutation (not present in chimp, gorilla, gibbon or macaque) was also present in the Altai Neanderthal. There was one difference between the Neanderthal and human *CLUU1* sequences. The amino acid at position 110, Lysine (K), is encoded by AAG in Neanderthal and by AAA in human. The presence of the enabler and lack of disruptive mutations suggest that Neanderthal possessed a functional *CLUU1* gene. Surprisingly, given the sister group relationship between Neanderthal and Denisovan, the Denisovan did not possess the enabler ΔA. Thirty-six reads covered the position and all showed the adenine. Denisovan also had the ancestral AAG triplet that encoded a Lysine in Neanderthal. These three nucleotides cannot be considered a codon in the Denisovan, nor in chimp and macaque. Without the upstream ΔA, a TGA (stop) disrupts the ORF long before this triplet in all three species

Neanderthals and Denisovans appear to have functional *DNAH10OS* and *C22orf45* genes. At the *DNAH10OS* locus both species possess the ‘enabling’ 10 bp insertion and neither have mutations that disrupt the *DNAH10OS* ORF, though they did share two derived substitutions (synapomorphies): They had a synonymous, third position T to G substitution in codon 23: CC**G** in Neanderthal and Denisovan vs CC**T** in human, and a nonsynonymous G to A substitution in codon 153: **A**GT in Neanderthal and Denisovan vs **G**GT in human. This second change causes a Glycine to Serine amino acid substitution. The human, Neanderthal and Denisovan *C22orf45* gene sequences were identical.

These analyses show that the enabling mutations necessary for the ‘origination’ of all three de novo genes occurred at least 800,000 years ago in the common ancestor of Neanderthals, Denisovans and modern *Homo sapiens*. While this conclusion does not help establish function, it does show that, whatever they do, *CLUU1*, *C22orf45*, and *DNAH10OS* are unlikely to be responsible for human-specific traits. The *CLUU1* data suggest that the coding and non-coding alleles persisted in the common ancestor of Neanderthals and Denisovans for several hundred thousand years, which indicates a lack of strong selection on either allele. This observation also suggests that genetic drift was less operative in this ancestor (*H. heidelbergensis*) than it appears to have been in its descendants.

